# Apparent growth paradox of *Staphylococcus epidermidis* in response to extremely low frequency electromagnetic radiation exposure

**DOI:** 10.1101/2021.03.06.434218

**Authors:** Dyaan Malik, Esmeralda Pineda, Deyvis Mejia Zambrana

## Abstract

*Staphylococcus epidermidis* is a normal part of the human microbiome; however, it is an opportunistic pathogen and can cause infections when the delicate balance of this microbiome is disrupted. Furthermore, infections caused by this bacterium can be hard to treat as a result of antibiotic resistance and biofilm production. This experiment aimed to determine whether electromagnetic field radiation (ELF-EMF) could be a deterrent of bacterial growth, as an alternative treatment to antibiotics. A non-pathogenic strain of *S. epidermidis* was used for experimentation, which took place in a school laboratory setting. The experimental group was exposed to ELF-EMF, while the control group did not receive the ELF-EMF treatment. The number of bacterial colonies, represented as colony forming units (CFUs) and area of random colonies were calculated to determine the effect of this treatment. There was no dramatic difference of colony formation on days 0, 1, and 2 of the four day period of ELF-EMF exposure. However, colony formation for days 3 and 4 showed a significant difference between the control and the experimental groups, as the experimental group had a significantly higher CFU count than the control. The average CFU count for day 3 in the control group was 420.6 and 1,097.4 for the experimental group (p<0.0001, t=12.9803). On the final day of the experimentation (Day 4) the average CFU count for the control group was 424.6 and 896.4 for the experimental group (p<0.0001, t=5.8926). The area for five randomly chosen colonies from each petri dish was calculated on the fourth day of experimentation. The area for the experimental group was significantly lower than that control (p<0.0001, with t=6.8659). The average area for the control group was 1.3249 mm^2^ and a lower average of 0.6375mm^2^ for the experimental group. These results demonstrate that the ELF-EMF treatment had an inhibitory effect on the area growth of *S. epidermidis*, but not on the colony-forming ability of S. epidermidis. This suggests that ELF-EMF influences the means by which the bacterium *S. epidermidis* grows.

## Introduction

The rapid surge of antibiotic and drug-resistant bacteria has been recognized as a serious problem worldwide; the infections and diseases that result from such bacteria jeopardize the lives of millions of people each year. Random mutations in bacteria allow for immunity to drugs that would once have been able to eliminate these bacteria. In other words, these bacteria gain resistance to antibiotics and survive and reproduce- increasing the presence of this trait in the population. Many people are unaware of these antibiotic-resistant bacteria and unintentionally promote their expanding population by misusing and overusing antibiotics. The unnecessary use of antibiotics increases the bacteria’s exposure to them and in turn augments the development of drug-resistant bacteria, putting more people at risk. Over 2 million people in the United States become infected with antibiotic-resistant bacteria each year and at least 23,000 of those people die as a direct result of these bacteria (CDC, 2013).

The lack of proper sterilization in hospitals, which causes infections to spread more rapidly, and the deficit of new drugs and vaccines are also linked to the rise of drug-resistant bacteria. A high percentage of hospital-acquired infections are caused by bacteria resistant to drugs which increases the rates of mortality in health care facilities (Struelens *et al.*, 1998). An alternative treatment needs to be developed that will kill antibiotic-resistant strains of bacteria so that infections or illness caused by these bacteria can be cured.

Infections caused by antibiotic-resistant bacteria can develop into life-threatening diseases if left untreated, which in many cases is what occurs due to the lack of alternative treatments for antibiotic-resistant bacteria. This experiment aimed to determine the effect of ELF EMF radiation on the growth of the bacterial colonies formed by *Staphylococcus epidermidis*. It was hypothesized that if *S. epidermidis* was exposed to an extremely low frequency electromagnetic field (ELF-EMF) then the viable count of colony forming units (CFUs) would decrease.

*S.epidermidis* is a bacterium that is very common in the world and in the environment around humans. *S. epidermidis* needs to be attached to a host and is commonly found in the normal human flora. The bacterium may be hazardous as it is an opportunistic pathogen; it attacks once the host has a weakened immune system (Brown *et al.*, 1988). Although *S. epidermidis* is usually not harmful, it has been known for causing many difficult-to-treat infections. *S. epidermidis* is notorious for causing infections in hospitals, specifically in prosthetic heart valves, prosthetic joints, and large wounds, due to its ability to form adhesive structures on medical devices such as catheters (Christensen *et al.*, 1982). *S. epidermidis* is also known for causing prevalent nosocomial infections which are hospital-acquired infections; these include urinary tract infections, surgical site infections, bloodstream infections, and pneumonia (Gill *et al.*, 2005).

An extremely low frequency electromagnetic field (ELF-EMF) was used to decrease the CFU count of *S. epidermidis*. Electromagnetic fields are a form of radiation that can change the chemical composition of organisms and physical objects (“Non-ionizing Radiation (EMFs)”, n.d., para. 1). EMFs were used due to the fact that there have been studies that have shown that certain types of EMFs have decreased the growth of cells (Zimmerman, *et al.*, 2013; Khaki, *et al.*, 2016). Zimmerman’s research suggests that the decrease in cell growth when exposed to ELF-EMF is caused by the cellular damage to internal structures that impedes the proliferation of cells. In Zimmerman’s study which was performed on human tumor cells, it was recorded that exposure to EMFs causes the size of tumors to decrease by interfering with the cell replication process (Zimmerman *et al.*, 2013). In Khaki’s study, cytotoxic effects of ELF-EMF demonstrated to be detrimental to ovarian follicles, oocytes, gametogenesis, and gonadal tissue. For treatment purposes an EMF-pump-mediated ELF-EMF was emitted through a device called an EMF pump in which the pump emits a field with ELF type frequencies. To take proper measures of protection against possible EMF hazards, aluminum cylinders were used to contain electromagnetic energy (Weibler *et al.*, 1993).

The effect of the EMF was measured by counting the bacterial colonies or colony forming units (CFU) formed by *S. epidermidis* on each Petri dish. CFU is an estimate of the cells that are able to grow under experimental conditions (Sutton *et al.*, 2011 A). Since the bacteria would be cultivated in petri dishes, the CFU count is appropriate to use as the low frequency EMF treatment will be performed over limited growth time and the bacteria will have limited space to populate. The formation of CFUs aids in the identification of a viable cell count for bacteria. One CFU, of a specific bacterial inoculation, represents one colony; a greater quantity of colony forming units counted indicates that the bacteria were able to proliferate in the presence of the ELF-EMF treatment. Thus, CFU count would reveal the efficacy of the treatment; a greater CFU count indicates cells were able to reproduce more easily than cells with a lower CFU count. The CFU counting method would allow for measuring growth between the higher and lower parameters of viable CFU count, these may be between 30 and 300 CFU (Sutton *et al.*, 2006). CFU counts below the minimum countable threshold would be negligible and labeled as too little to count, CFU counts greater than 300 are labeled as too numerous to count (TNTC).

The research methods for EMF exposure were adapted from previous research for *S. epidermidis* so that new treatment for antibiotic resistant bacteria would be explored where resistance is not acquired. The various methods for counting plated bacteria that grew from a stock solution include using a spectrophotometer to measure cell growth indirectly and counting colonies in CFU on a plate manually (Sutton *et al.*, 2011 A; Sutton *et al.*, 2011 B). The spectrophotometer calculates the density of cells by transmitting a light through a filter, then a detector calculates how much light was absorbed (passed through) a suspension containing cells. The CFU method was used instead of the optical density method as unnecessary exposure to any other type of radiation such as visible light may result in inadequate or skewed data measurements.

Methods for growing bacteria include a suspension and inoculation on agar plates (Aslanimehr *et al.*, 2013). Bacteria were inoculated on agar plates because colonies would be easier to count on a petri dish. Inoculation can be performed using an inoculator, however a unique method was developed which involved glass beads to spread a homogenous mixture over the surface of a petri dish. Thus, serial dilutions were necessary in applying a uniform amount and concentration of inoculum to both experimental and control groups. These quantitative methods were chosen as this proved to be more meticulous in terms of attention to detail; more variables would be greatly controlled, so that data may be most accurate to the greatest extent with the methods used.

## Methods & Materials

This experiment tested whether exposure to ELF-EMF would decrease the growth of *S. epidermidis* measured in CFU over 4 days and area on the 4th day. According to European and U.S. bacterial colony count standards, the countable CFU range must be between 30 and 300 so that it is feasible and capable of properly counting. The colony area was used as a measure that described colony growth and bacterial growth indirectly.

To begin inoculation, 0.5mL of *S. epidermidis* inoculum was added to 4.5mL of dH_2_O. Serial dilutions were then performed 5 times as shown in figure 4. Using a sterile micropipette, 100μL of 0.001% from original stock was transported and ejected into all 10 prepared agar plates for inoculation. Miniature spherical glass beads were then used to spread the solution immediately onto the surface of the solid agar. Strips of parafilm were used to further isolate an environment for bacterial colonies to grow and reduce environmental contaminants that would influence bacterial growth. All surfaces and materials used were sterilized with 4% bleach after experimentation and gloves were used when handling bacteria and disinfectants.

**Figure 1a.**
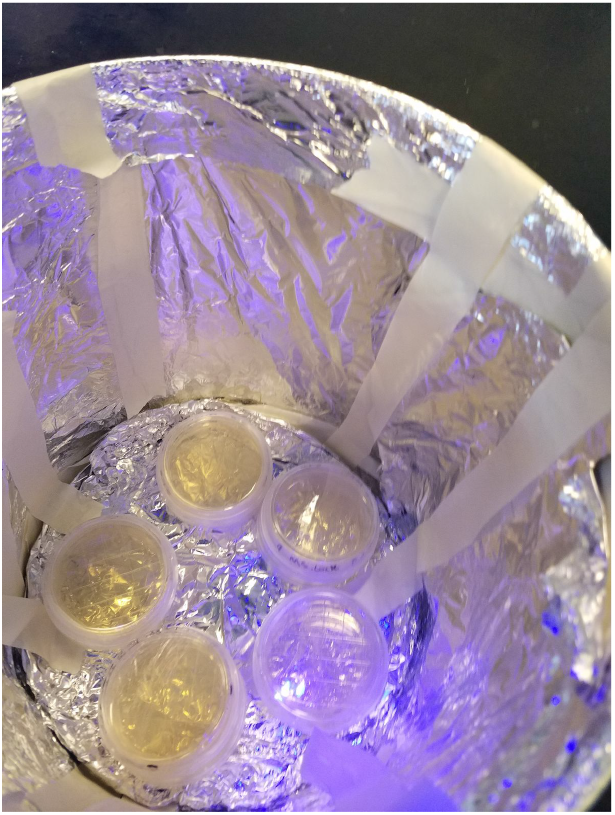
Petri dishes were stacked and rotated daily inside the aluminum cylinders.

**Figure 1b.**
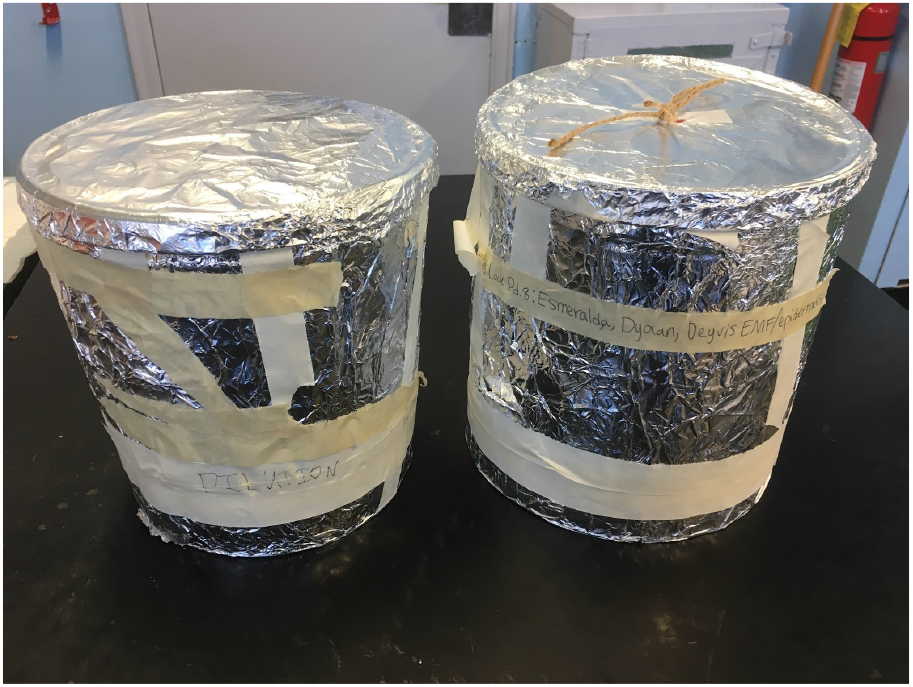
Aluminum cylinders were set up in the environment covered in aluminum.

**Figure 2.**
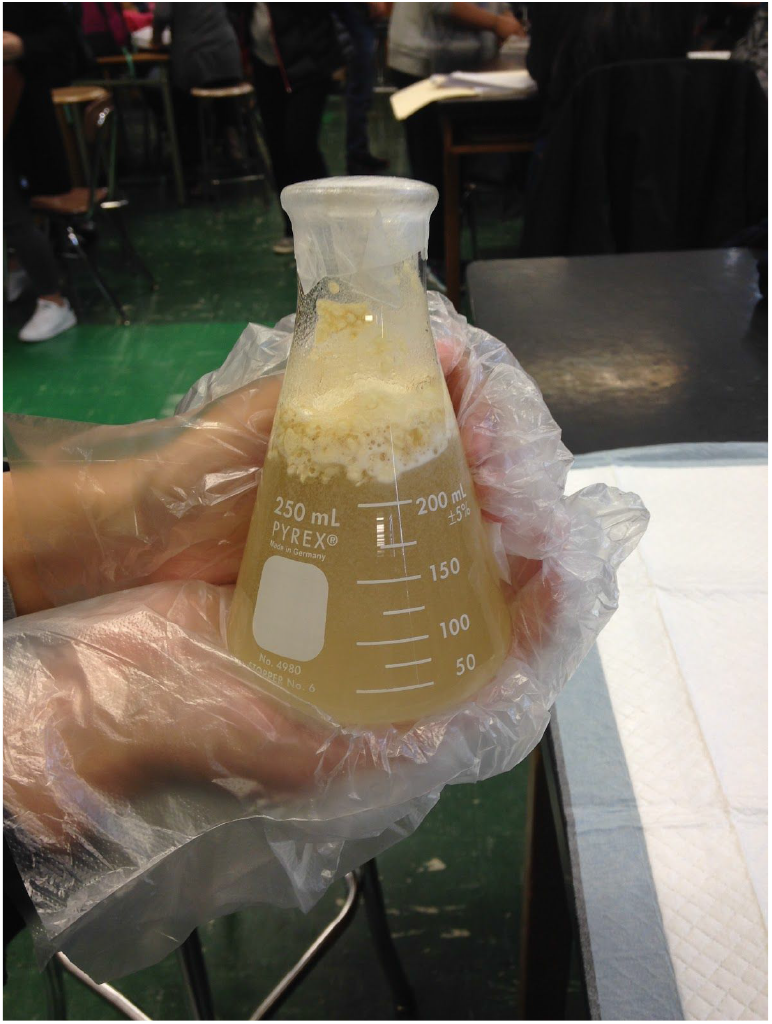
The image below shows the luria bertani agar before it was autoclaved and poured into petri dishes.

**Figure 3.**
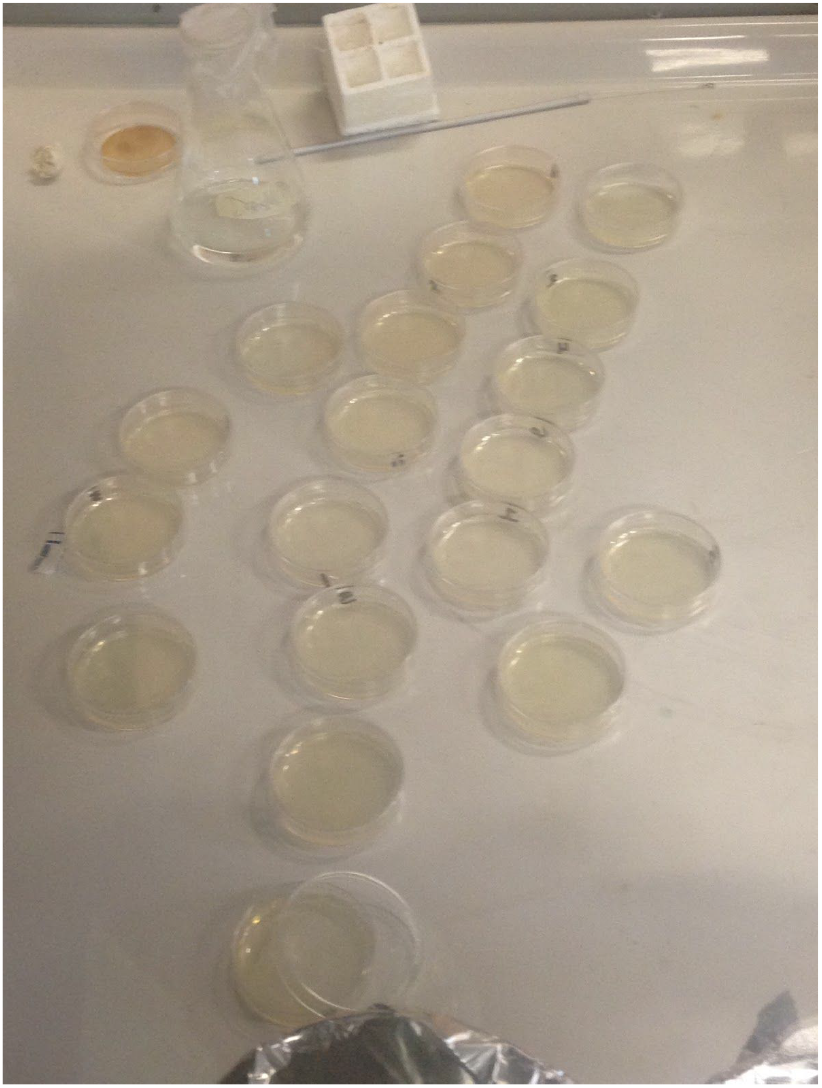
illustrates the agar on petri dishes kept under the fumigation hood to limit contamination.

**Figure 4.**
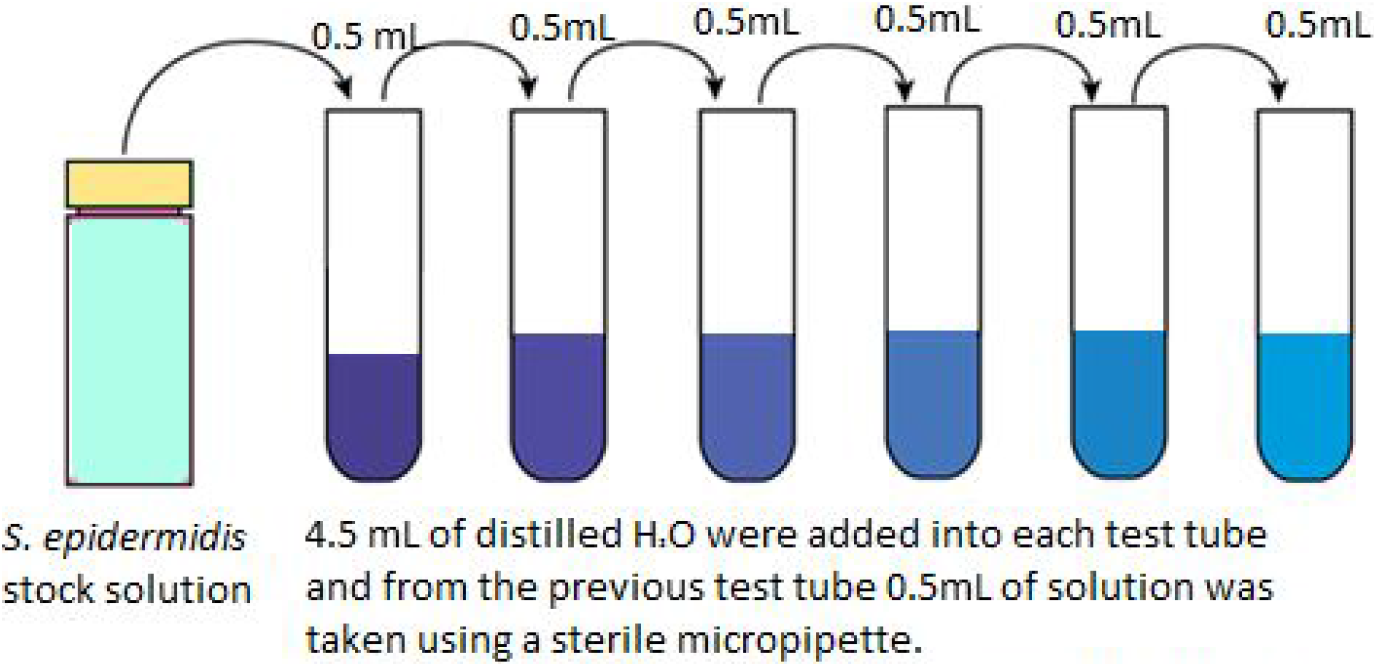
Serial dilution procedure.

The circular petri dishes used for inoculation were 15×60 mm in dimension. Luria bertani (LB) agar was used as a general medium to cultivate *S. epidermidis*. 6.4g of agar were mixed with 200mL of water using a magnetic stir plate (figure 2) and autoclaved at 250° C for one hour to limit contamination of the agar. After the agar cooled down for 20 minutes, it was poured into 20 petri dishes about halfway. These were then placed under a fumigation hood, previously sterilized with 4% bleach, for 10 minutes to prevent the buildup and condensation of excess water in the petri dishes (figure 3).

Separate aluminum cylinders were used to isolate the groups as shown in figures 1a and 1b since aluminum blocks out any external EMF radiation and can retain EMF radiation that is emitted from the EMF pump inside of the experimental cylinder (Weibler, *et al.*, 1993). The cylinder used for the control group only contained 10 inoculated and sealed Petri dishes. The cylinder used for the experimental group contained the EMF pump and 10 Petri dishes (also inoculated and sealed) evenly spaced out around the EMF pump. The EMF pump that was inside the experimental cylinder was turned on and the EMF frequency was measured using an EMF meter placed on the inside of the cylinder. It was assumed that the EMF meter indicated that the battery-operated EMF pump was producing an electromagnetic field at a frequency of 15 hertz as indicated by the distributor. The EMF pump was kept on for 4 days for about 20 hours each day of experimentation.

After 4 days, the mean data represented how many colonies were formed by *S. epidermidis* on average for the experimental and control groups. 10 random colonies were selected from each petri dish for both experimental and control groups on the fourth day. A dissecting microscope and computer software (Amscope) were used to measure the diameter of each randomly chosen colony which was then used in calculating the areas and finally averaging the areas.

## Results

The data were collected every day over a period of four days. In total there were 20 petri dishes inoculated, including the experimental and control groups, with a non pathogenic strain of *S. epidermidis*. The number of colony forming units (CFUs), which measures cell viability for which a single cell would produce a colony, was measured over the four day period. On the fourth day of data collection, area was also used to measure the growth, specifically the spread of the colonies that arose in both groups. The raw data were organized into a table for quantitative analysis. Measures of central tendency were used to summarize the data sets for the control and experimental groups. The measure of central tendency that best represented the raw data overall was the arithmetic mean, shown in tables 3 and 4.

**Table 3:** Average Area for Colony Growth for Experimental and Control Groups.

| Control (without exposure to EMF) | Average Area (mm<sup>2</sup>) | Experimental (with exposure to EMF) | Average Area (mm<sup>2</sup>) |
| --- | --- | --- | --- |
| Petri dish #1 | 1.089 | Petri dish #1 | 0.490 |
| Petri dish #2 | 1.020 | Petri dish #2 | 0.653 |
| Petri dish #3 | 1.500 | Petri dish #3 | 0.658 |
| Petri dish #4 | 1.262 | Petri dish #4 | 0.641 |
| Petri dish #5 | 1.266 | Petri dish #5 | 0.656 |
| Petri dish #6 | 1.311 | Petri dish #6 | 0.720 |
| Petri dish #7 | 0.916 | Petri dish #7 | 0.548 |
| Petri dish #8 | 2.000 | Petri dish #8 | 0.578 |
| Petri dish #9 | 1.517 | Petri dish #9 | 0.726 |
| Petri dish #10 | 1.368 | Petri dish #10 | 0.705 |
| **Total Average** | **1.3249** | **Total Average** | **0.6375** |

**Table 4:**
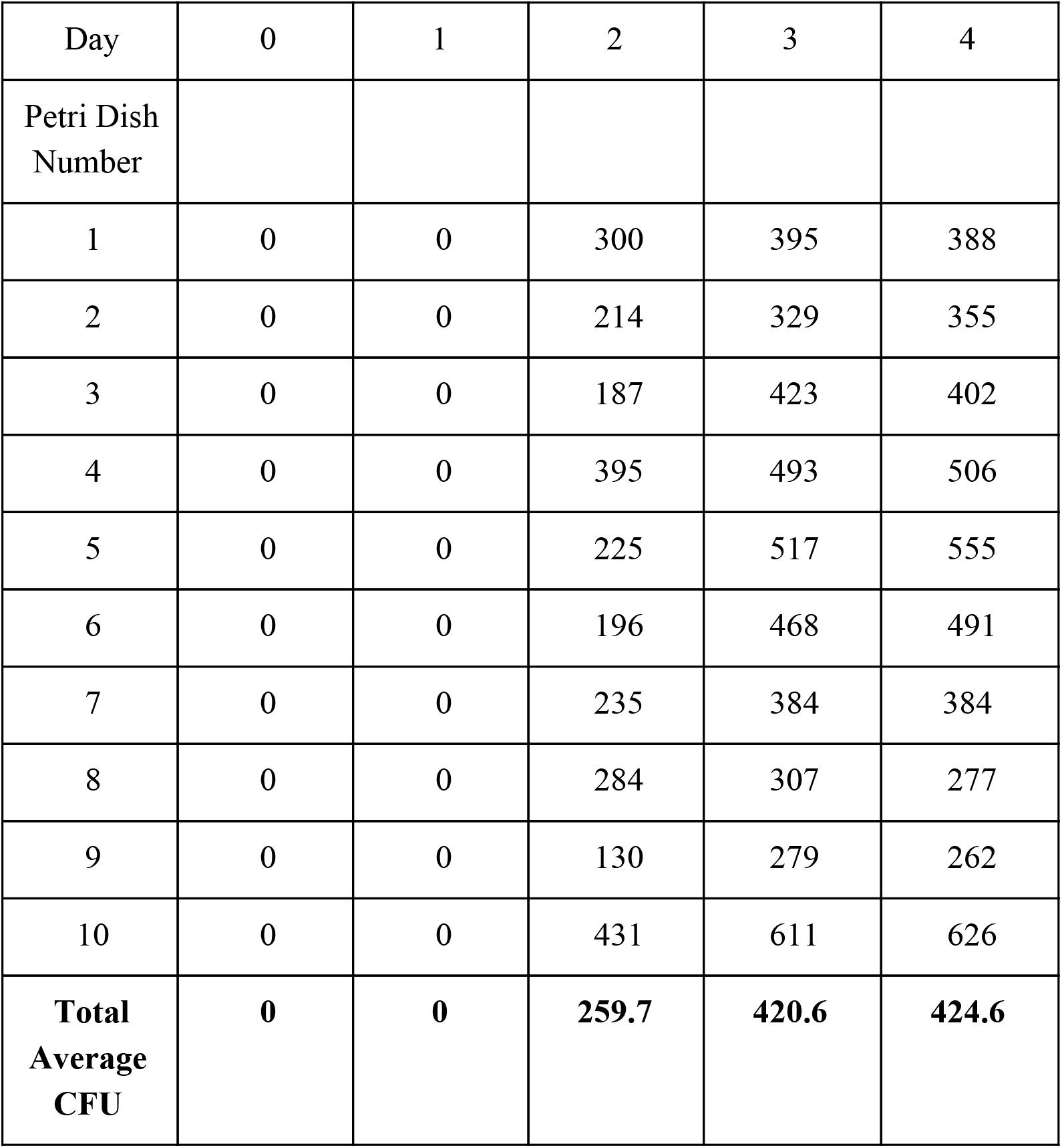
Average CFU Counts of 10 Petri Dishes Over a Four Day Period for the Control Group. The average CFU counts for the control group were averaged for all Petri dishes for every day as shown above. Lower CFU counts on consecutive days suggest is attributed to human error. There is overall positive growth (CFU counts) for the control group over time.

| Day | 0 | 1 | 2 | 3 | 4 |
| --- | --- | --- | --- | --- | --- |
| Petri Dish Number |  |  |  |  |  |
| 1 | 0 | 0 | 300 | 395 | 388 |
| 2 | 0 | 0 | 214 | 329 | 355 |
| 3 | 0 | 0 | 187 | 423 | 402 |
| 4 | 0 | 0 | 395 | 493 | 506 |
| 5 | 0 | 0 | 225 | 517 | 555 |
| 6 | 0 | 0 | 196 | 468 | 491 |
| 7 | 0 | 0 | 235 | 384 | 384 |
| 8 | 0 | 0 | 284 | 307 | 277 |
| 9 | 0 | 0 | 130 | 279 | 262 |
| 10 | 0 | 0 | 431 | 611 | 626 |
| **Total Average CFU** | **0** | **0** | **259.7** | **420.6** | **424.6** |

### Part 1: Area

Figure 5 and table 3 show that the average area for colony growth was higher in the control group than in the experimental group. The average area for 100 randomly chosen colonies from the control group was 1.3249 mm^2^, while the average area for the 100 colonies randomly selected from the experimental group was 0.6375 mm^2^. After running the data collected from the average area for colony growth, a t test was performed on this data, the results are extremely statistically significant (p < 0.0001 and t=6.8659). The p value of p<0.0001 demonstrates that the mean for the two groups which are the Control (no treatment) and Experimental (with treatment) are significantly different; this suggests a causal relationship between treatment and measurement.

**Figure 5.**
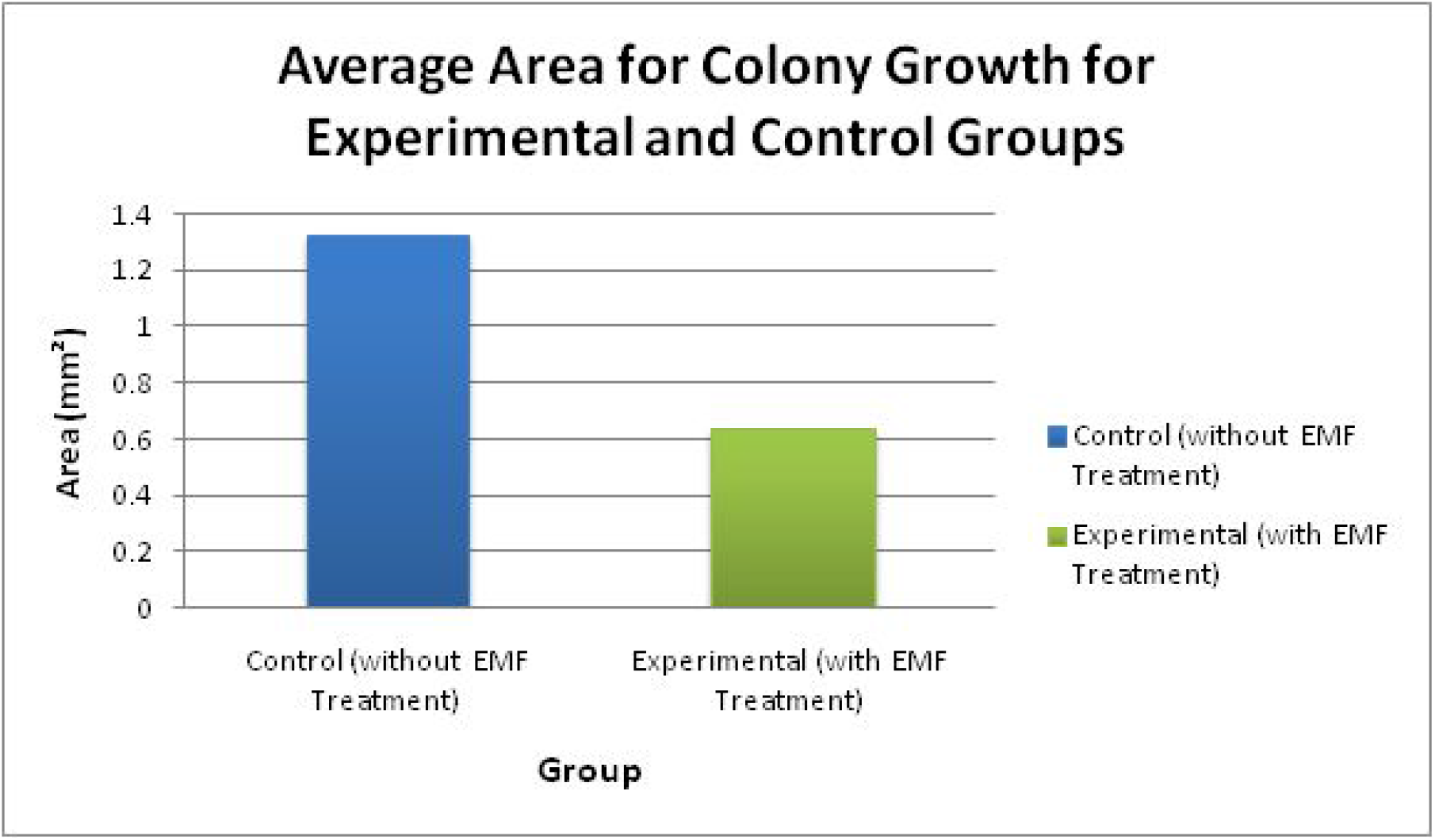
Colony area measured on the final day oftreatment exposure. The graph depicts the average size of a colony for each group. The control group demonstrated colonies that were larger in size than compared to the experimental group. The asterisk above the experimental group indicates that the value for the experimental group is statistically significant.

The table above demonstrates the average size of the colonies that grew for both the control group and the experimental group on the fourth day of data collection. Data was collected by using Amscope, a computer program, to select 5 random colonies from each petri dish and calculate the diameter of each random colony. Then the radii of each colony was calculated to solve for the area of each colony. Afterwards, the area of the 5 colonies in each petri dish was averaged so that the average area for that petri dish was recorded. After that for each group the average area for each of the 10 petri dishes within the group was averaged to have a final average area for the whole group. Based on the data it is shown that the control group tended to grow colonies that were larger than those of the experimental. The data is extremely statistically significant, the P value is less than 0.0001 and t=6.8659 for this set of data.

### Part 2: CFU

Figure 6 and tables 4, 5, and 6 demonstrate that the average CFU count was higher in the experimental group than in the control group. This is statistically significant because p<0.0001 and t=5.8926 for day 4; p<0.0001 and t=12.9803 for day 3. This significance shows a causal relationship between the EMF treatment and the lower CFU counts. On days 3 and 4 there were greater CFU counts in the experimental group than the control groups. Day 3 had an average of 420.6 CFU for the control group and an average of 1,097.4 for the experimental group; on day 4 there was an average of 424.6 CFU for the control group and an average of 896.4 CFU for the experimental group (table 6).

**Table 5:** Average CFU Counts of 10 Petri Dishes Over a Four Day Period for the Experimental Group. The average CFU counts for the experimental group were averaged for all Petri dishes for every day as shown above. Lower CFU counts on consecutive days is attributed to human error. There is overall positive growth (CFU count) for the experimental group over time.

| Day | 0 | 1 | 2 | 3 | 4 |
| --- | --- | --- | --- | --- | --- |
| Petri Dish Number |  |  |  |  |  |
| 1 | 0 | 0 | 491 | 1,270 | 646 |
| 2 | 0 | 0 | 391 | 1,198 | 560 |
| 3 | 0 | 0 | 634 | 1,094 | 701 |
| 4 | 0 | 0 | 579 | 958 | 902 |
| 5 | 0 | 0 | 503 | 1,110 | 1,027 |
| 6 | 0 | 0 | 504 | 1,026 | 939 |
| 7 | 0 | 0 | 559 | 1,075 | 998 |
| 8 | 0 | 1 | 406 | 800 | 757 |
| 9 | 0 | 0 | 424 | 1,049 | 1,062 |
| 10 | 0 | 0 | 782 | 1,394 | 1,372 |
| **Total Average CFU** | **0** | **0.1** | **413.5** | **1,097.4** | **896.4** |

**Table 6:** Average CFU Counts Over a Four Day Period. The table demonstrates that throughout the 4 day period the experimental group had more growth than that of the control group. Specifically on Days 3 and 4 there was a greater amount of growth in the experimental than the control.

| Day | 0 | 1 | 2 | 3 | 4 |
| --- | --- | --- | --- | --- | --- |
| Control Group<br>(without EMF<br>exposure) Average<br>CFU | 0 | 0 | 259.7 | 420.6 | 424.6 |
| Experimental (with<br>EMF exposure)<br>Average CFU | 0 | 0.1 | 413.5 | 1,097.4 | 896.4 |

**Figure 6.**
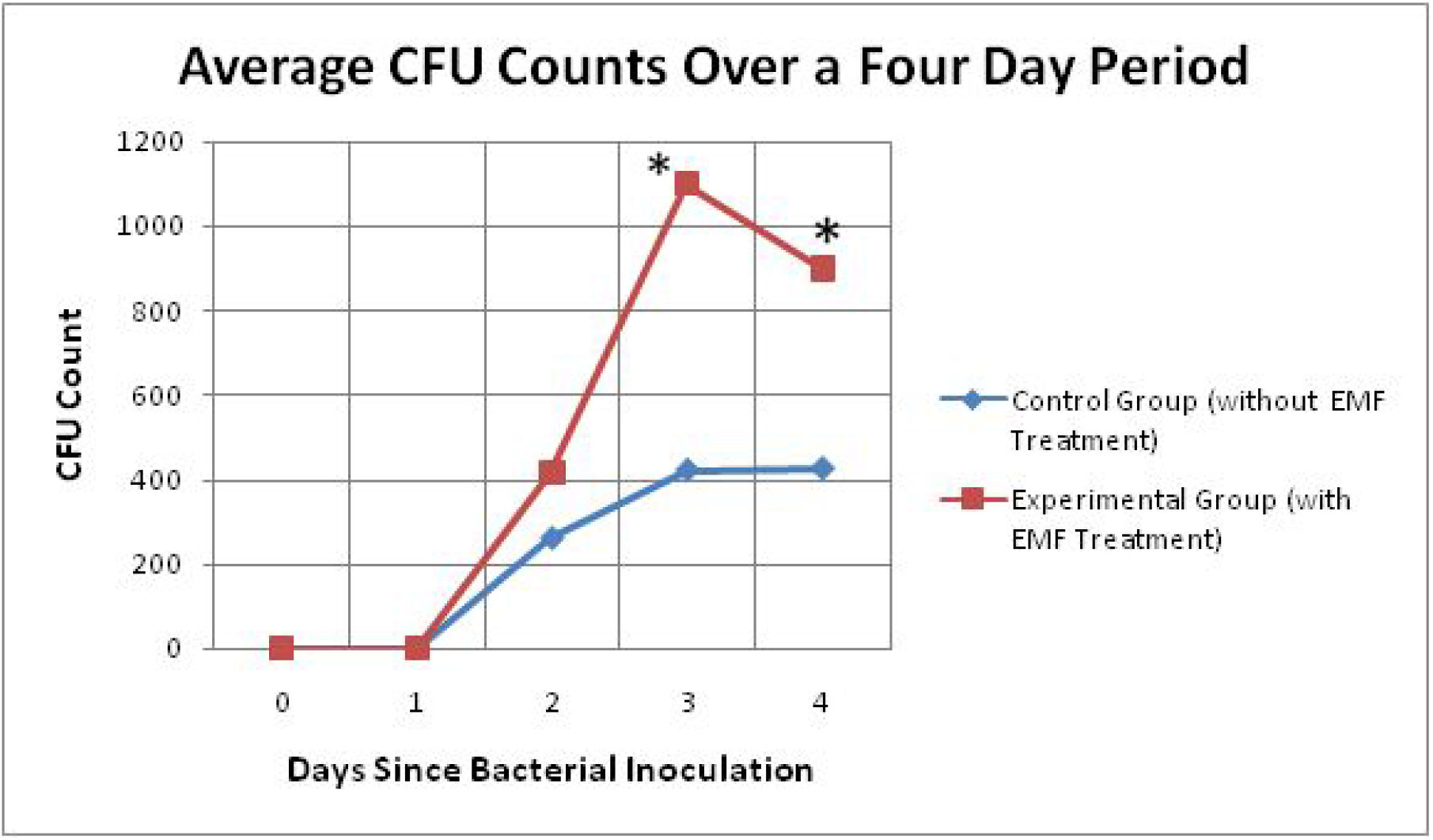
CFU counts over a four day treatment period. The graph shows the trend between the experimental and control group for a four day period. It depicts the average of 10 petri dishes from each group for each day. The trends demonstrate that throughout the 4 day period the experimental group had more growth than the control. Also as shown Days 3 and 4 have an asterisk above the data for the experimental group for that day. The asterisk indicates that the experimental data is significant and that the results are not due to chance for days three and four.

## Discussion

The hypothesis was partially supported by the data analyzed because the area was significantly lower in the experimental group than in the control group, Although the CFU count for the experimental group on days 3 and 4 were significantly higher than that of the control group, the results suggest that there is a controlled mechanism within *S. epidermidis* that links the way it spreads over a surface and the viability of S. epidermidis to give rise to a colony. Non-ionizing ELF-EMF radiation affects the average area in growth of bacterial colonies as well as the number of colonies formed. The results of overall increase in CFU counts suggests that cell division is stimulated by exposure to ELF-EMF.

Research suggests that the decrease in cell growth when exposed to ELF-EMF is caused by the cellular damage to internal structures that impedes the proliferation of cells. In a study performed on human tumor cells, it was recorded that exposure to EMFs causes the size of the tumor to decrease (Zimmerman *et al.*, 2013). The decrease in tumor size suggests that the frequencies of the EMF disrupted the process of cell division. Similar to this study, the experiment conducted demonstrated a decrease in size of each bacterial colony. However, contrary to the study, the experiment demonstrated no decrease in CFU.

Although the hypothesis was partially supported, limitations indicate that the results could be products of chance as the sample size was small, so data would have been skewed to a greater degree than if a larger sample size was used, in addition to the averaging of all the data collected. These limitations include the strain of *S. epidermidis*, refrigeration time of the inoculum, unseen biological contamination, varying levels of oxygen, high humidity in the petri dishes, temperature fluctuations, and vibrations from the EMF pump. Biological contamination may have affected the growth if there was competition for the resources in the petri dishes, and the unavailability for the usage of organic molecules present. Some fungi grew in petri dishes which therefore needed to be remade. When pouring the agar onto the petri dishes there may have been contamination from the exhaled human breath which could have provided the microorganisms associated with the human respiratory system a way of colonizing and competing in the petri dish with *S. epidermidis*. Research suggested that the competition of bacteria in the human nasal cavity is highly competitive (Lina *et al.*, 2003). Research also discusses that cellular products called bacteriocins may give bacteria a competitive edge (Riley *et al.*, 1999). The growth of bacteria might have been the result of vibrations emitted by the EMF pump since the attachment of extracellular cell structures onto surfaces anchor the cell for replication to occur (Christensen *et al.*, 1982). In addition, the strain of *S. epidermidis* was unknown which may have affected the ability for the bacteria to grow in sufficient quantities to be countable.

By using only two measurements of growth, limitations could have developed since other methods of measuring growth may have allowed for more quantitative data to describe how *S. epidermidis* grows as well as better insight into the connection between ELF-EMF exposure and the growth of *S. epidermidis*. Area described the ability to spread in a growth medium of an *S. epidermidis* colony. This measurement may have been influenced by other factors such as human errors while handling the bacteria and petri dishes, as well as temperature differences between the experimental and control groups. Human errors throughout this experiment include the counting of colony-forming units, recording data, and incorrectly inoculating the agar with bacteria. Temperatures may have fluctuated differently in both groups due to the fact that temperature could not be controlled in the school laboratory in which the experiment took place.

A study has suggested that high humidity may deter microbial growth (Loosdrecht *et al.*, 1987). If there were different levels of humidity in each petri dish then growth may have varied due to inhibition of growth by preventing the attachment of cells onto a surface, especially if the cell membrane becomes increasingly hydrophobic as they do at high growth rates. Refrigeration of prolonged time may have slowed down the metabolism of *S. epidermidis*, thus prolonging the lag phase of bacterial growth and resulting in lower CFU counts. A study conducted proposed that the reproducibility of experiments involving bacterial stock solution that were in cold storage could be valid in their reproducibility for at least four days (Zaharia *et al.*, 2010). In addition to this, it is known that temperature has a great impact on the ability for cells to multiply, higher temperatures would have resulted in higher CFU counts, however an incubator was not used as it was too small for the closed systems to fit. The experiment was conducted at around room temperature, so the inner closed system remained relatively cool for the most part.

The applications of this research remain strong in the area of antibiotic resistance; however, further experimentation with a larger sample size is needed to establish a concrete trend within the observed data. The effects of ELF-EMF exposure on cell growth are not generally known. This experiment may prove to be valuable for general information purposes that can be applied into the setting of the physical world and health since ELF-EMF is a non-ionizing type of radiation, which means that it is located at the low end of the electromagnetic spectrum and is less dangerous to humans than ionizing radiation (WHO, n.d.). Limitations will be addressed, respectively, to improve the experiment by using a more concentrated serial dilution to yield higher colony counts.There will also be more trials to further diminish the impact contamination has on the process of data collection and analyzing for patterns. Other bacteria will also be used to verify that this method of combating bacteria is a viable one that could be applied to other bacteria that are antibiotic resistant. Questions that arose pertain to how changes in conditions internal or external to *S. epidermidis* change the way it reproduces.

## Acknowledgements

We would like to thank Mrs. Lock and Mrs. Homer, our research teachers from these past few years. Ms. Kim, the AP of Science. Mr. Pedicini and Mr. Obaji for letting us use their facility as well as their resources.New York Institute of Technology (NYIT) for a research grant allowing us to continue conducting our research. We feel very grateful for all of the help and support.

## Author Contributions

All authors were involved in experimental design, data collection, analysis, background research and manuscript drafting.

